# Signatures of optimal codon usage predict metabolic ecology in budding yeasts

**DOI:** 10.1101/2020.07.22.214635

**Authors:** Abigail Leavitt LaBella, Dana A. Opulente, Jacob Steenwyk, Chris Todd Hittinger, Antonis Rokas

## Abstract

Reverse ecology is the inference of ecological information from patterns of genomic variation. One rich, heretofore underutilized, source of ecologically-relevant genomic information is codon optimality or adaptation. Bias toward codons that match the tRNA pool is robustly associated with high gene expression in diverse organisms, suggesting that codon optimization could be used in a reverse ecology framework to identify highly expressed, ecologically relevant genes. To test this hypothesis, we examined the relationship between optimal codon usage in the classic galactose metabolism (*GAL*) pathway and known ecological niches for 329 species of budding yeasts, a diverse subphylum of fungi. We find that optimal codon usage in the *GAL* pathway is positively correlated with quantitative growth on galactose, suggesting that *GAL* codon optimization reflects increased capacity to grow on galactose. Optimal codon usage in the *GAL* pathway is also positively correlated with human-associated ecological niches in yeasts of the CUG-Ser1 clade and with dairy-associated ecological niches in the family Saccharomycetaceae. For example, optimal codon usage of *GAL* genes is greater than 85% of all genes in the major human pathogen *Candida albicans* (CUG-Ser1 clade) and greater than 75% of genes in the dairy yeast *Kluyveromyces lactis* (family Saccharomycetaceae). We further find a correlation between optimization in the thiamine biosynthesis and *GAL* pathways. As a result, optimal codon usage in thiamine biosynthesis genes is also associated with dairy ecological niches in Saccharomycetaceae, which may reflect competition with co-occurring microbes for extracellular thiamine. This work highlights the potential of codon optimization as a tool for gaining insights into the metabolic ecology of microbial eukaryotes. Doing so may be especially illuminating for studying fungal dark matter—species that have yet to be cultured in the lab or have only been identified by genomic material.

## INTRODUCTION

The immense diversity of life is due, in part, to adaptation to the wide variety of environmental niches available. By acting on the interface between genotype, phenotype, and environment, natural selection has given rise to numerous ecological adaptations [1–3]. The precise relationship between genotype, phenotype, and environment, however, is often elusive. For example, a connection was only recently made between environmental distribution of seeds of different sizes, phenotypic variation in the beaks of Darwin’s finches, and changes in the expression of the protein BMP4 [4–6].

Genomic sequencing has accelerated the rate at which the underlying genomic mechanisms of well-established ecologically adapted phenotypes are elucidated [7,8]. While powerful, this type of ecological genomics requires extensive knowledge of the ecological niche in which species live. For many microbial species, however, detailed ecological information is unavailable due to both the scale of the ecosystems they live in and the dearth of information reported during collection [9]. One potentially powerful way to address this gap in knowledge is to use the extensive genomic resources available in microbes to conduct reverse ecology – directly inferring ecology from genotype [10,11].

Reverse ecology has successfully linked environmental phenotype with genotype using multiple types of genomic features [11–13]. Optimal growth temperature was successfully inferred from genomic content, including tRNA, ribosome, and gene features, in 549 Bacteria and 170 Archaea [14]. In the red bread mold *Neurospora crassa*, analysis of highly divergent genomic regions in 48 isolates uncovered “genomic islands” associated with adaptation in two different ecosystems [15]. Across the entire tree of life, metabolic capability (assessed using Kyoto Encyclopedia of Genes and Genomes (KEGG) gene annotations) was used to examine the evolution of exogenously required metabolites likely found in the environment [16]. Metabolic network analysis has emerged as a common genomic feature for reverse ecology analysis [17,18]. There are, however, other promising genomic features that can be used in reverse ecology.

One underutilized genomic feature with great potential for reverse ecology studies is codon usage, which has long been associated with gene expression [19–21]. Changes in gene expression have been shown to play an important role in ecological adaptation [22–24]. For example, in wild isolates of budding yeast *Saccharomyces cerevisiae*, changes in the expression of multiple genes were associated with phenotypic differences in copper resistance and pigmentation that may be associated with high copper environments [25]. Over evolutionary time, increased levels of gene expression result in a selective pressure for accurate and efficient translation [26–30] and increased mRNA stability [31,32]. Codons that match the tRNA pool— called optimal codons—have a substantial impact on both translation [27,29,30] and mRNA stability [31]. Therefore, optimal codon usage is correlated with high gene expression in multiple lineages, especially in microbes [19,33–38]. Therefore, we hypothesize that ecological adaptations that are, at least partly, due to high expression levels of specific genes or pathways will be reflected in their codon usage values.

Previous work in diverse microbes supports the hypothesis that codon optimization can be used to identify associations between codon usage (either globally or in specific genes) and ecology [12,39–43]. For example, an analysis of metagenomes collected from mine biofilms shows an enrichment of optimal codons in bacterial and archaeal genes associated with inorganic ion transport[39]. In fungi, codon optimization in host-induced and secreted proteins is associated with generalist fungal parasites [41]. Although these studies were highly successful in linking particular ecological niches with highly enriched groups of genes, we still lack examples where reverse ecology has linked particular ecologies to specific pathways.

The galactose (*GAL*) pathway (also known as the Leloir pathway) in the budding yeast subphylum Saccharomycotina is an iconic pathway that metabolizes galactose into glucose, which can then be used in core metabolism or as an intermediate [44,45]. The genes encoding the three enzymes of the *GAL* pathway—*GAL1* (encoding a galactokinase), *GAL10* (encoding a UDP-glucose-4-epimerase), and *GAL7* (encoding a galactose-1-phosphate uridyl transferase)*—* are frequently clustered in yeast genomes and are induced in response to the presence of galactose [46–48]. There has been extensive research into the biochemistry [44], regulation [49–51], and evolutionary history [48,52] of this pathway. Ecological work on the *GAL* pathway has revealed that gene inactivation is associated with an ecological shift in *Saccharomyces kudriavzevii*, a close relative of the species to *S. cerevisiae* [53]. There is also a positive association between galactose metabolism ability and the flower/*Ipomoea* isolation environment and a negative association between galactose metabolism ability and tree or insect frass isolation environments [54]. While gene gain and loss in budding yeasts may play an important role in ecological adaptation, variation in gene expression is also a likely contributor [55–57]. The recent publication of 332 budding yeast genomes and the identification of translational selection on codon usage in a majority of these species provide a unique opportunity to test for differences in *GAL* gene expression—inferred from optimal codon usage—across ecological niches inferred from recorded isolation environments [54,58–60].

In this study, we characterize the presence and codon optimization of the *GAL* pathway in 329 budding yeast species and identify an association between optimization in the *GAL* pathway and two specific ecological niches. We identify a complete set of *GAL* genes in 210 species and evidence of physical clustering of *GAL1, GAL7*, and *GAL10* in 150 species. Consistent with our hypothesis that codon optimization is a signature of high gene expression, we find that growth rate on galactose-containing medium is positively and significantly correlated with *GAL* codon optimization. In the CUG-Ser1 major clade, which contains the opportunistic human pathogen *Candida albicans*, codon optimization in the *GAL* pathway is higher in species found in human-associated ecological niches when compared to species associated with insect (and not human) ecological niches. In the family Saccharomycetaceae, another major clade in the Saccharomycotina subphylum, which contains the model species *S. cerevisiae*, we find that codon optimization in the *GAL* pathway is higher in species isolated from dairy-associated niches compared to those from alcohol-associated niches. For example, codon optimization among closely related *Kluyveromyces* species is nearly twice as high in species isolated from dairy niches as those found associated with marine or fly niches. We also used KEGG Orthology (KO) annotations to find metabolic pathways with codon optimization that correlated with *GAL* optimization. We identified multiple members of the thiamine biosynthesis pathway whose codon optimization is not only correlated with galactose metabolism, but associated with specific ecological niches. This study serves as a foundation for future high-throughput reverse ecology work that uses codon optimization to link metabolic pathways with ecological niches in microbes.

## METHODS

### Galactose (*GAL*) Pathway Characterization

Genomic sequence and gene annotation data were obtained from the comparative analysis of 332 budding yeast genomes [58] (Supplementary Table 1). Mitochondrial sequences were filtered from these genomes using previously described methods[59]. Reference protein sequences for *GAL* gene annotation (approximately 40 proteins for each of the *GAL* genes) were obtained from GenBank and previous KEGG ortholog (KO) annotations [58,61]. A protein HMM profile was constructed for each *GAL* gene and used to conduct two HMMER searches (version 3.1b2; http://hmmer.org/), one on publicly available annotations and one on all possible open reading frames generated using ORFfinder (version 0.4.3; https://www.ncbi.nlm.nih.gov/orffinder/). The search on all possible open reading frames was done to ensure that inferences of *GAL* gene absences were not due to errors in publicly available gene annotations. The results of the two searches were compared using the Perl script fasta_uniqueseqs.pl (version 1.0; https://www.ncbi.nlm.nih.gov/CBBresearch/Spouge/html_ncbi/html/fasta/uniqueseq.cgi). Discrepancies between the two searches, which most often occurred in cases where the publicly available annotation combined two nearby genes, were resolved manually. The genes *GAL1* and *GAL3* are known ohnologs (i.e., paralogs that arose from a whole genome duplication event) [62,63]. Thus, the identity of *GAL1* and *GAL3* genes was inferred for *Saccharomyces* species by phylogenetic analysis of the *GAL1/3* gene tree constructed using the IQ-Tree webserver (http://iqtree.cibiv.univie.ac.at/; default parameters; Supplementary Figure 1)[64–66]. Other *GAL1* homologs were included as there is a lack of evidence for functional divergence in other lineages [51]. All reference and annotated *GAL* genes are available in the supplementary FigShare repository. All instances where *GAL1, GAL7*, and *GAL10* were found on the same contig were considered to represent *GAL* gene clusters.

**Figure 1:**
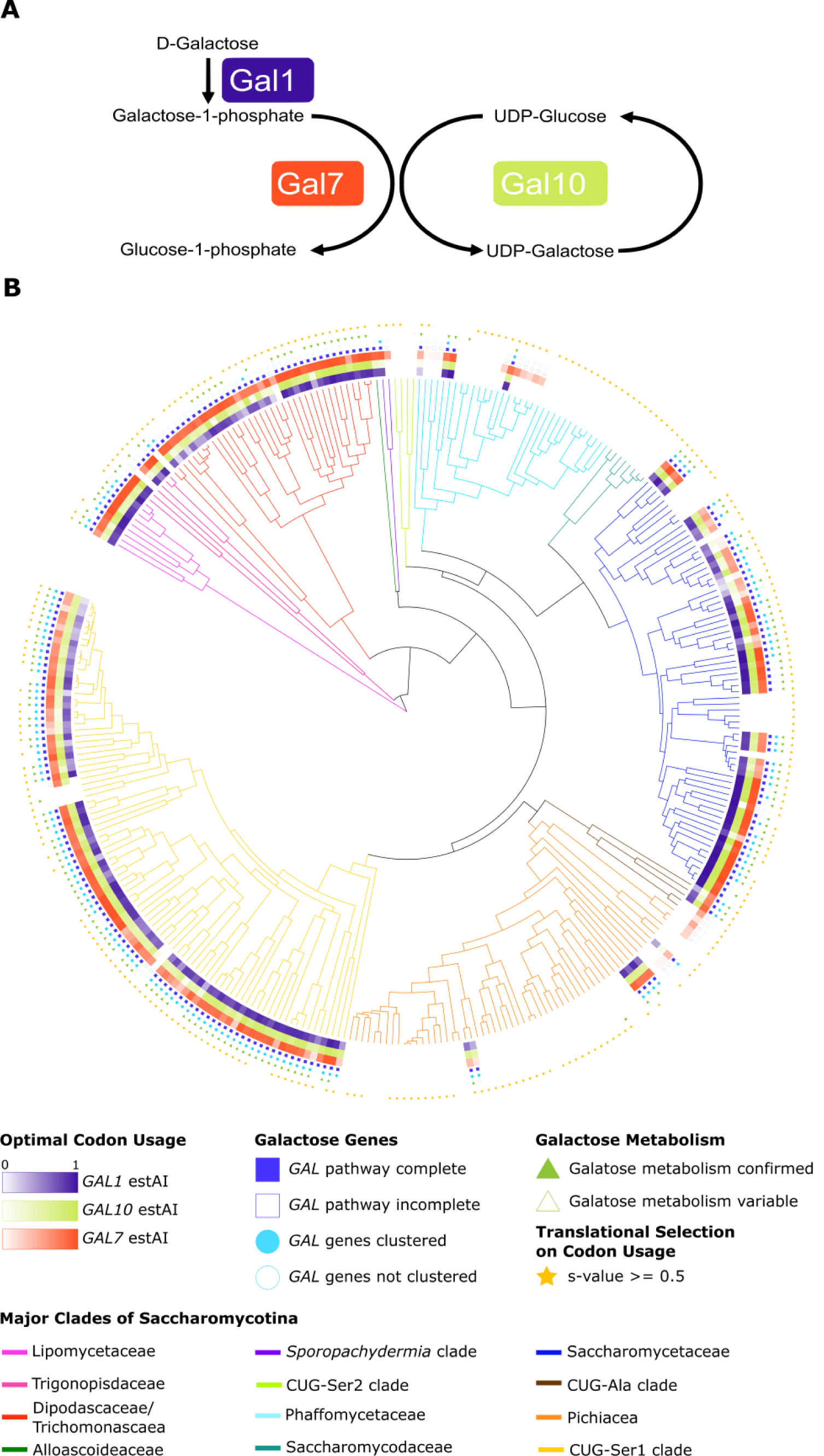
The *GAL* pathway and the distribution of galactose metabolism, *GAL* genes, and preferred codon usage across the Saccharomycotina. A) The three enzymes of the *GAL* pathway metabolize galactose into glucose-1-phosphate, which can then enter glycolysis after being converted into glucose-6-phosphate. B) Various features of galactose metabolism plotted on a phylogeny of the budding yeast subphylum Saccharomycotina; the 12 major clades of the subphylum are color-coded. The presence and codon optimization (measured by estAI) of the three *GAL* genes are represented in the inner three rings. We did not identify any *GAL* genes from species in the CUG-Ser2 clade or the family Saccharomycodaceae. High codon optimization (darker colors) in the *GAL* pathway is not restricted to any one major clade. Complete and clustered occurrences of the *GAL* pathway (filled-in blue squares and circles respectively) are found in every other major clade examined. The ability to metabolize galactose (filled-in green triangle) was assessed either experimentally in this study or taken from the literature. In some instances where only literature data were available, there were conflicting or variable reports of galactose metabolism (5 species; empty green triangles). The majority of species in the Saccharomycotina have also been shown to have genome-wide selection on codon usage (denoted by the yellow stars)[59].

### Codon Optimization in the *GAL* pathway

To infer gene expression in the *GAL* pathway, we calculated the level of codon optimization in each *GAL* gene and compared it to the genome-wide distribution of codon optimization. Codon optimization of individual *GAL* genes was assessed by calculating the species-specific tRNA adaptation index (stAI) from previously calculated species-specific codon relative adaptiveness (wi) values [59,67]. Three species that previously failed to generate reliable wi values (*Martiniozyma abiesophila, Nadsonia fulvescens* var. *elongata*, and *Botryozyma nematodophila*)[59] were removed from all subsequent analyses. The stAI software does not take into account the CUG codon reassignment in the CUG-Ser1 and CUG-Ala clades. Previous analysis, however, suggests that this codon is rare [59] – the average frequency of the CUG codon in species where it has been reassigned is 0.005, 0.003, and 0.006 for *GAL1, GAL10*, and *GAL7*, respectively – and its influence on codon optimization calculations is not significant.

The stAI for each gene was calculated by taking the geometric mean of the wi values for all the codons, except the start codon. The genome-wide distribution of gene stAI values is normally distributed, but the mean varies between species [59]. To compare codon optimization between species, we normalized each gene’s stAI value using the empirical cumulative distribution function to get the percentage of all genes with stAI values lower than that of the gene of interest; we call this the estAI value. For example, an estAI value of 0.4 for a given gene would indicate that 40% of the genes in the genome have lower stAI values (i.e., are less optimized) than the gene of interest. The estAI optimization values therefore range from 0 to 1, with 1 being the most optimized gene in the genome.

A total of 49 species’ genomes had multiple copies of at least one *GAL* gene. For those genomes, the gene with the highest estAI value was used. For example, we identified two copies of *GAL10* in *Candida ponderosae* located on different contigs with estAI values of 0.46 and 0.44. Therefore, we used the estAI value of 0.46 as the representative *GAL10* value for this species. The average difference between the maximum and minimum estAI for multiple copies of *GAL1, GAL7*, and *GAL10* are 0.0948, 0.0007, and 0.0125. There were 14 cases where all gene copies with the highest estAI values were not found on the same contig. In 18 cases, all duplicates with the highest estAI values were located on the same contig. The use of the *GAL* gene copy with the highest estAI is supported by evidence in *S. cerevisiae* that functionally derived gene duplicates have reduced codon optimization, which is likely linked to an evolutionary trajectory towards novel functions [68].

### Galactose Growth Data

To test the hypothesis that high levels of *GAL* codon optimization are associated with strong growth in media where galactose is the sole carbon source, we measured galactose and glucose (as a control) growth for 258 species in the laboratory. Yeast strains corresponding to the species whose genomes were sequenced were obtained from the USDA Agricultural Research Service (ARS) NRRL Culture Collection in Peoria, Illinois, USA or from the Fungal Biodiversity Centre (CBS) Collection in the Netherlands. All strains were initially plated from freezer stocks on yeast extract peptone dextrose (YPD) plates and grown for single colonies. YPD plates were stored at 4°C until the end of the experiment. To quantify growth on galactose and glucose, we set up three replicates on separate weeks using different colonies for each strain. Strains were inoculated into liquid YPD and grown for six days at room temperature. For each replicate, strains were randomized and arrayed into a 96-well plate. The plate was then used to inoculate strains into a minimal medium containing 1% D-galactose or 1% glucose, 5g/L ammonium sulfate, and 1.7g/L Yeast Nitrogen Base (w/o amino acids, ammonium sulfate, or carbon) and grown for seven days. After a week, we transferred all strains to a second 96-well plate containing fresh minimal medium containing galactose or glucose.

To quantify the growth of each strain/species, we measured its optical density (OD units at 600nm) following growth in a well of a BMG LABTECH FLUOstar Omega plate reader after a week at room temperature. We calculated two measures of growth, growth rate and endpoint, for each species and replicate. The growth rates were calculated in R (x64 3.5.2) using the grofit package (v 1.1.1.1) and end point, a proxy for saturation, was calculated by subtracting the T_0_ time point from the final time point for each species. We visually assessed growth on galactose for all species using the growth curves we collected; a species was denoted as having the ability to grow on galactose if it grew in at least 2 of 3 replicates tested. Growth data, both growth rate and endpoint, were set to zero for all species that did not meet this requirement. Quantitative growth on galactose was successfully measured for a total of 258 species. Growth on galactose was then computed relative to glucose to account for differences in the baseline growth rate of different species due to variables, such as cell size and budding type (unipolar versus bipolar).

For the 71 species where new quantitative galactose growth data were unavailable, we used previously published species-specific binary growth data [54,58,60]. Uncertain growth is indicated where conflicting or variable growth was found in the literature (empty green triangles; Figure 1B). Quantitative galactose growth data (normalized to glucose) were compared to maximum gene codon optimization values using phylogenetically independent contrasts (PIC)[69]. Data from related species are not independent observations and therefore require a PIC analysis to ensure that covariation between traits is not the result of the relatedness of species [69]. The PIC analysis was conducted in R using the ade4 package [70]. The species *Metschnikowia matae* var. *matae* was removed from this analysis as it was a clear outlier on the residual plots for a complementary PGLS analysis (Supplementary Figure 2)[71,72]. Outliers in phylogenetically independent analyses occur when two closely related taxa have disparate trait values, which can be identified by examining the residual plots. In this case, the taxa *Metschnikowia matae* var. *maris* (yHMPu5000040795 / NRRL Y-63737 / CBS 13985) and *Metschnikowia matae* var. *matae* (yHMPu5000040940 / NRRL Y-63736 / CBS 13986) are very closely related, and yet the growth rate on galactose for *Metschnikowia matae* var. *matae* (1.390) is nearly double that of *Metschnikowia matae* var. *maris* (0.750) and the next most closely related species *Metschnikowia lockheadii* (0.567).

**Figure 2:**
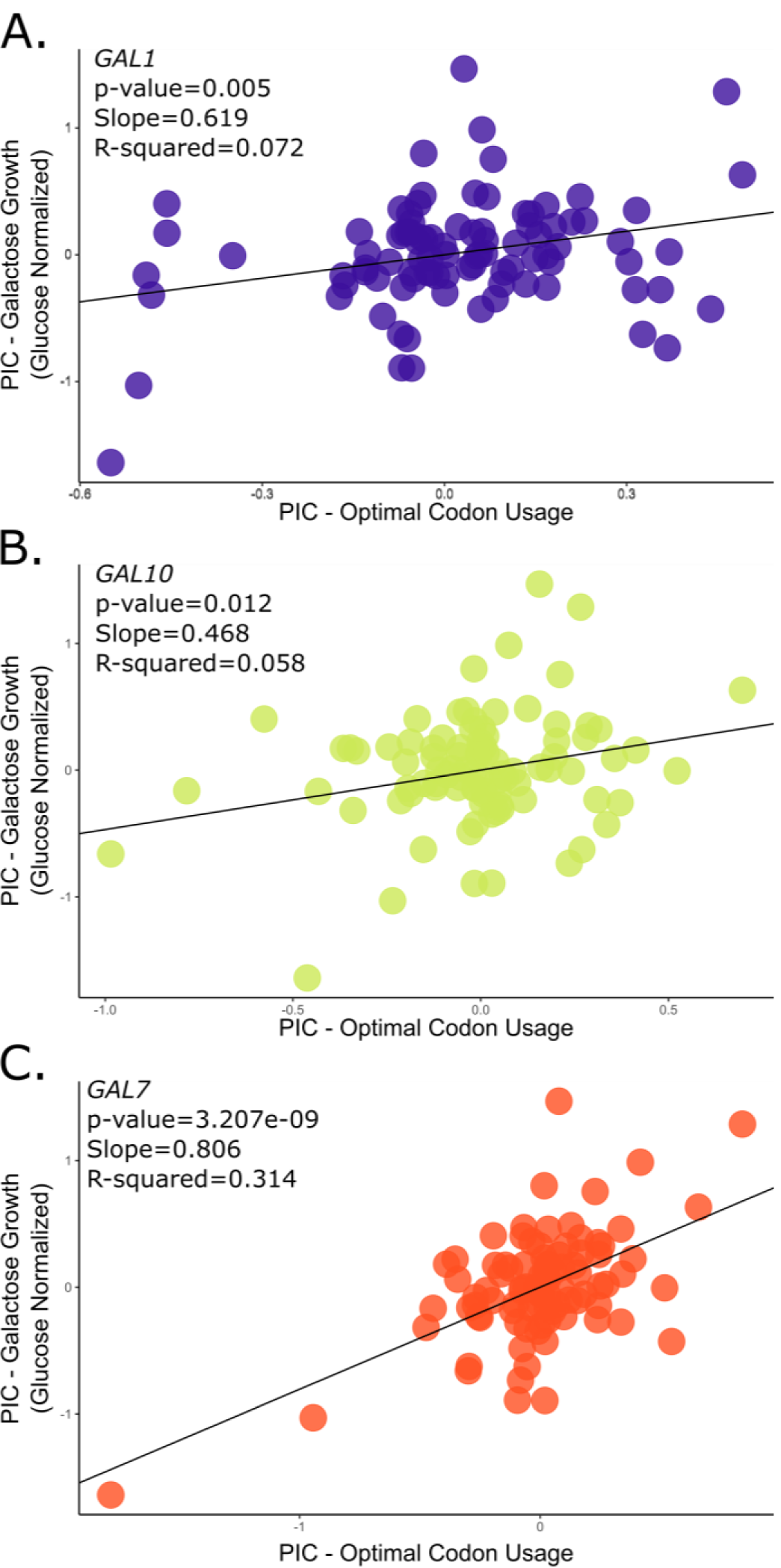
Codon optimization in the *GAL* pathway is positively and significantly correlated with growth rate on galactose. Phylogenetically independent contrasts (PIC) analyses of galactose growth (Y axis) versus *GAL* gene optimal codon usage (X axis). There is a significant and positive correlation between the PIC values for codon optimization and galactose growth in *GAL1* (A), *GAL10* (B), and *GAL7* (C). The best fit and strongest correlation is between growth on galactose and optimization in *GAL7* (C). The analyses included 94 species with a growth rate on galactose greater than 0, a complete *GAL* cluster, and evidence of genome-wide translational selection on codon usage. One species, *Metschnikowia matae* var. *matae*, was removed as an obvious outlier based on residual analysis.

### Ecological association analysis

To test for associations between *GAL* pathway codon optimization and ecological niche, we obtained species-specific isolation data from multiple sources. We first tested 50 isolation environments from data collated from *The Yeasts: A Taxonomic Study*[58,60], as recorded by Opulente and coworkers [54,58]. We compared codon optimization in each of the *GAL* genes between species isolated from a given environment versus species not isolated from that environment (Supplementary Figure 3). From this analysis, we identified four general ecological niches with potentially differential codon optimization: dairy-, alcohol-, insect-, and human-associated ecological niches. To validate and update the data from *The Yeasts*, we conducted an in-depth literature search for these four specific ecological niches for each of the 329 species of interest using all known anamorphs and synonyms per species (see Supplementary Table 2 for updated information for the ecological niches and associated references). Dairy ecological niches identified included milks, butters, cheeses, and yogurts. Alcohol ecological niches identified included spontaneous beer fermentation, alcohol starters, wine, ciders, kombuchas, and liquors. Insect-associated ecological niches included insect guts, insect bodies, and insect frass. Human-associated ecological niches were characterized as any isolation from a human, regardless of pathogenicity. Additionally, we did not take into account studies where species identification lacked genetic data and relied solely on phenotypic and assimilation data, because these identifications have been shown to be potentially unreliable [73–75]. For example, the only evidence that the species *Candida castellii* is associated with dairy niches comes from a single identification in a fermented milk product using only metabolic chacterization [76]. Therefore, *C. castellii* was not considered associated with dairy niches.

**Figure 3:**
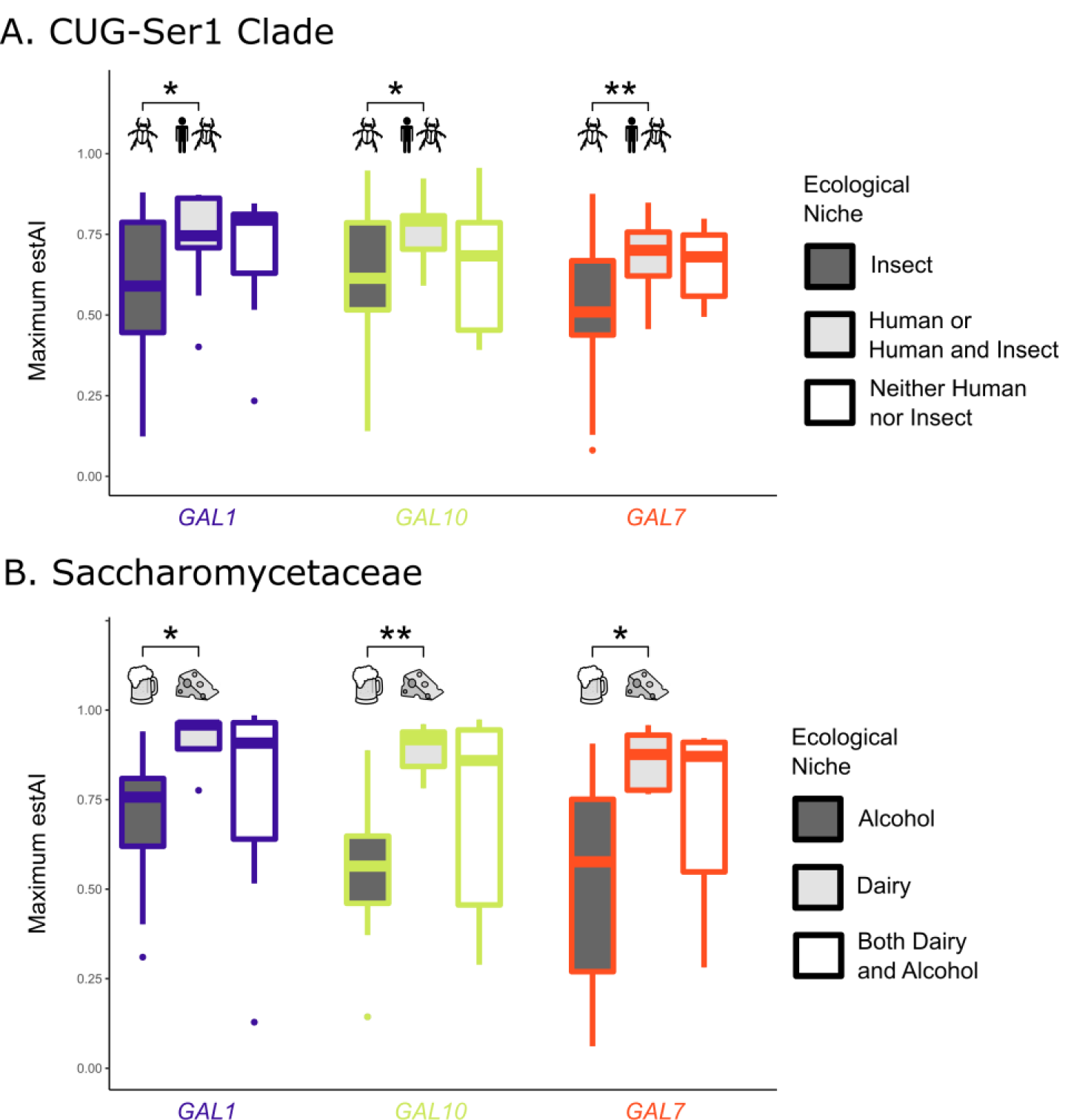
Codon optimization in the *GAL* pathway is correlated with specific ecological niches in two different major clades of budding yeasts. P-values less than 0.01 are indicated with ** and less than 0.05 with *. A) In the CUG-Ser1 clade, species associated with a human niche or human and insect niches (13 species) have significantly higher codon usage optimization values in all *GAL* genes (p-values of 0.022, 0.028, and 0.006 for *GAL1, GAL10*, and *GAL7*, respectively) when compared to species that are associated with insect niches but not human niches (44 species). Only 11 species were not associated with either human or insect niches. B) In the Saccharomycetaceae, species associated with only dairy niches (5 species) have significantly higher codon usage optimization values in all of the *GAL* genes (p-values of 0.010, 0.002, and 0.014 for *GAL1, GAL10*, and *GAL7*, respectively) versus species associated with only alcohol niches (14 species). A total of 9 species are associated with both dairy and alcohol niches.

To test for significant differences in *GAL* optimization between ecological niches, we first filtered the species set to retain only those that contain all three *GAL* genes (210 species) and that were previously shown to exhibit genome-wide selection on codon usage (266 species; s-value >=0.5)[59]; thus, the total number of species tested was 170. We then compared levels of *GAL* codon optimization between ecological niches using the Wilcoxon rank sum test in R [77].

### Evolutionary rate analysis

To examine variation in the evolutionary rates among *GAL* genes, we used the maximum likelihood software PAML (version 4.9)[78,79]. Specifically, we examined the rates of synonymous changes in the *Kluyveromyces* species using the free-ratios model that allows for a different rate of evolution along each branch. The species tree was used as the backbone tree, and nucleotide sequences were aligned using the codon aware software TranslatorX (http://translatorx.co.uk/)[80].

### Identification of additional metabolic pathways whose codon usage correlates with *GAL* optimization

To identify additional pathways that exhibit the same codon optimization trends between ecological niches as the *GAL* pathway, we tested whether the optimization of KEGG orthologs (KOs) was correlated with that of the *GAL* genes. KO annotations were previously generated for all species [58]. We started with the 266 genomes with evidence of translational selection on codon usage and identified 2,573 KOs present in 100 or more of those species. We then conducted a PIC analysis between the optimization of the *GAL* genes and each of the KOs across the species. P-values were adjusted to account for the total number of KOs tested using a Bonferroni correction (Supplementary Table 3). Based on the results of the PIC analysis, we further investigated the correlation between the thiamine biosynthesis pathway and the *GAL* pathway. To ensure we were not missing any members of the thiamine biosynthesis pathway, we annotated the entire pathway using the same method used for annotation of the *GAL* genes. We then re-ran the PIC analysis with the curated thiamine gene set.

## RESULTS & DISCUSSION

### Variable *GAL* pathway and codon optimization across the Saccharomycotina

To examine variation in *GAL* codon optimization across the subphylum, we first examined whether *GAL* genes were present in each of the 329 genomes. Across the Saccharomycotina, we annotated 742 *GAL* genes (265, 256, and 221 annotations for *GAL1, GAL10*, and *GAL7*, respectively) in a total of 233 species (Supplementary Table 1 and FigShare Repository). The complete *GAL* enzymatic pathway (i.e., *GAL1, GAL10*, and *GAL7*) was identified in 210 species, of which 149 had evidence of *GAL* gene clustering. We cannot, however, rule out clustering of the *GAL* genes in the remaining 61 species as some of the annotations were at the ends of the contigs.

There were some discrepancies between galactose growth data and *GAL* gene presence data. Three species where galactose growth was experimentally observed lacked all three *GAL* genes: *Ogataea methanolica, Wickerhamomyces* sp.YB-2243, and *Candida heveicola*. The growth rates for these species are 0.129, 0.339, and 0.211 for *O. methanolica, Wickerhamomyces* sp., and *C. heveicola*. The low growth rates (7^th^ and 3^rd^ lowest overall) of *O. methanolica* and *C. heveicola* suggest these species may be utilizing trace amounts of other nutrients present in the medium. Finally, there were 26 species with a complete *GAL* gene cluster where no growth on galactose has been reported. This may represent a loss of pathway induction in these species or an inability to induce growth in the specific experimental conditions tested, as observed previously in the genus *Lachancea* [81]. Inactivation of the *GAL* pathway has also occurred multiple times in budding yeasts [48,53], and some of these taxa could be in the early stages of pathway inactivation.

Codon optimization in the *GAL* pathway, measured by estAI, varied greatly across the Saccharomycotina (Figure 1B.) The estAI values ranged from 0.02 (or greater than only 2% of the genes in the genome) in *GAL7* from *Lachancea fantastica* nom. nud. to 0.99 (or greater than 99% of the genes in the genome) in *GAL1* from *Kazachstania bromeliacearum*. To determine if there is an association between codon optimization and the ability to grow on galactose, we compared optimization in the *GAL* pathway between species that are able and unable to grow on galactose. We found that species without evidence for growth on galactose had significantly lower (p < 0.05) codon optimization in *GAL1* and *GAL7* (Supplementary Figure 4). This correlation is consistent with a relaxation of selective pressures in non-functional pathways [82–84] and previous work has identified multiple parallel inactivation events of the *GAL* pathway in budding yeasts [53]. The *GAL* pathway may have alternative roles in cell function that are not associated with growth on galactose and may have not experienced the same selective pressures. For example, in *Candida albicans, GAL10* has been shown to be involved in cell integrity [85]. Finally, the *GAL* pathway may have an alternative induction system in these species. For example, the fission yeast *Schizosaccharomyces pombe* (not a member of the Saccharomycotina) has a complete *GAL* cluster but is unable to grow on galactose. Mutants of *S. pombe*, however, have been isolated that constitutively express the *GAL* genes and can grow on galactose [86].

**Figure 4:**
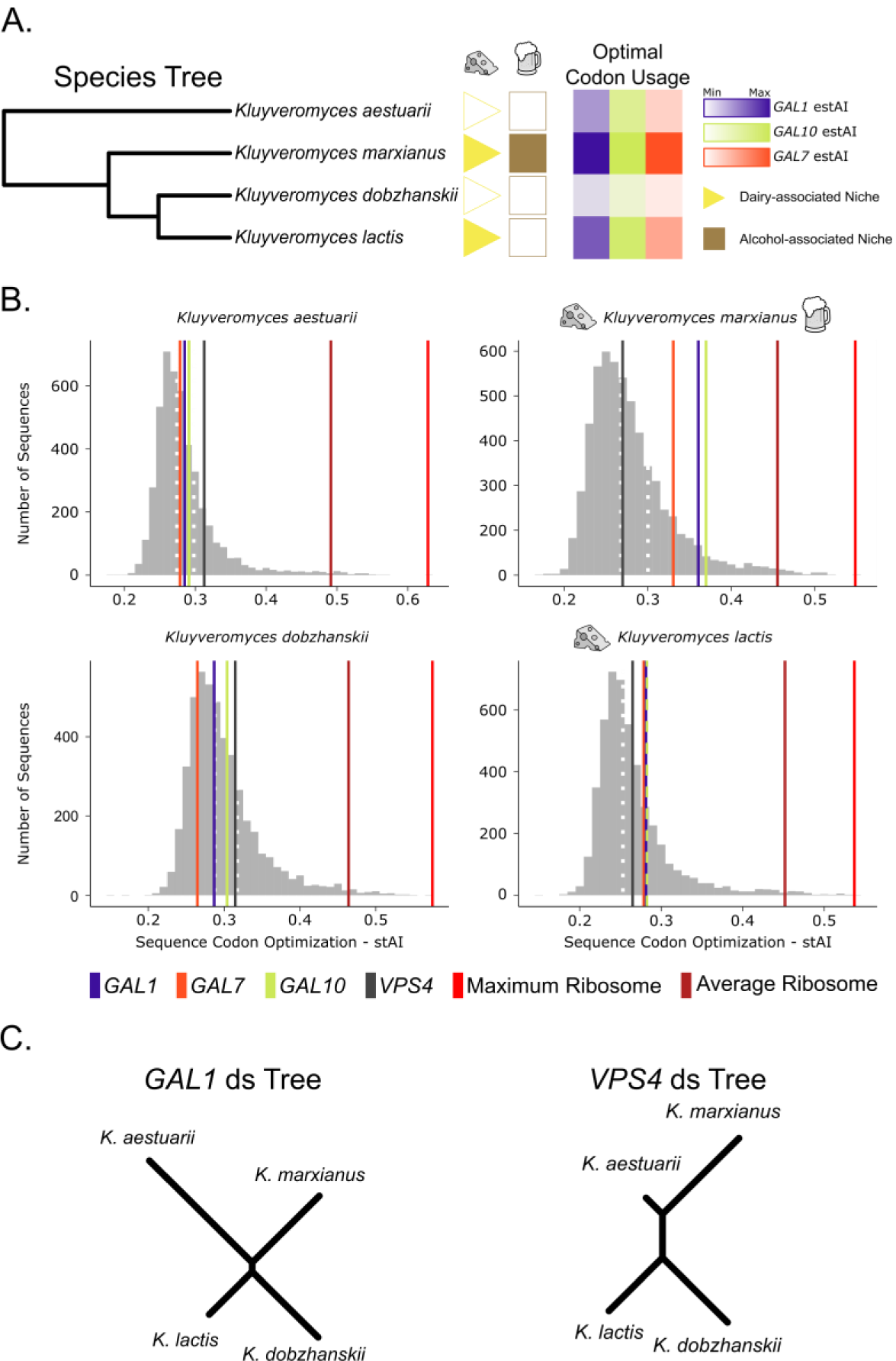
Closely related *Kluyveromyces* species exhibit differential codon optimization in the *GAL* pathway associated with isolation from dairy environments. All four *Kluyveromyces* species were shown experimentally to metabolize galactose. A) Species phylogeny of four closely related *Kluyveromyces* species. *K. marxianus* and *K. lactis* are both associated with dairy niches and have high codon optimization values in their *GAL* pathway genes. In contrast, *K. aestuarii* is associated with marine mud, and *K. dobzhanskii* is associated with flies. B) The genome-wide distribution of codon optimization (stAI) values for the four *Kluyveromyces* species included in this study. The 50^th^ and 75^th^ percentiles are shown with white dashed lines. In the two species associated with dairy niches, the codon optimization for all three *GAL* genes falls in the top 25^th^ percentile. In the two species not associated with dairy, the *GAL* genes fall below the top 25^th^ percentile. The gene *VPS4* (encoding a protein involved in <u>v</u>acuolar <u>p</u>rotein <u>s</u>orting) is a non-metabolic gene with intermediate codon optimization value across budding yeasts. Genes encoding ribosomal proteins are well established to rank among the most highly optimized genes within a genome. C) The unrooted trees show the estimated rate of synonymous substitutions in the *GAL1* and *VPS4* genes along these lineages. The long branch in *K. aestuarii* for the *GAL1* tree suggests a relaxation of selection on synonymous sites in this lineage.

### *GAL* codon optimization is correlated with growth rate on galactose

Strong translational selection on codon usage is correlated with highly expressed genes in diverse organisms [34,35,37,87–91]. Therefore, we hypothesized that high levels of codon optimization in the *GAL* pathway reflect high levels of *GAL* gene expression and ultimately high growth rates on galactose. To test this hypothesis, we measured growth rate on galactose relative to glucose. We found a significant positive correlation between growth rate on galactose-containing medium and codon optimization in the *GAL* pathway of genomes that have experienced translational selection on codon usage (N species = 94, linear regression of PIC values; p-values of 0.005, 0.012, and 3.207e^-9^ for *GAL1, GAL10*, and *GAL7*, respectively; Figure 2). Codon optimization of *GAL7* showed the strongest correlation with growth rate (Figure 2C), which may reflect the gene’s function; *GAL7* encodes for the enzyme that metabolizes galactose-1-phosphate, a toxic intermediate [92,93] whose accumulation has been shown to reduce growth rate in *S. cerevisiae* [93]. Furthermore, the correlation between *GAL7* optimization and growth rate on galactose remained strong when analyzed independently in both the Saccharomycetaceae (29 species) and in the CUG-Ser1 clade (47 species; Supplementary Figure 5), the two largest clades sampled. The *GAL1* and *GAL10* genes were both significantly positively associated with growth rate in galactose in the Saccharomycetaceae, but not in the CUG-Ser1 clade (Supplementary Figure 5). This contrast may reflect the different regulatory mechanisms involved in galactose assimilation in the two major clades—tight control via a regulatory switch in the Saccharomycetaceae versus leaky expression in CUG-Ser1 [49,50]. We also tested the correlation between growth rate on galactose containing medium and the *PGM*1 and PGM2 genes that encode phosphoglucomutases which converts\ glucose-1-phosphate (Figure 1) to glucose-6-phosphate. There was no correlation between optimization in *PGM1* or *2* and growth on galactose containing medium (Supplementary Figure 6). Collectively, our findings support the hypothesis that codon optimization is the result of selection on codon usage in species with high *GAL* gene expression.

**Figure 5:**
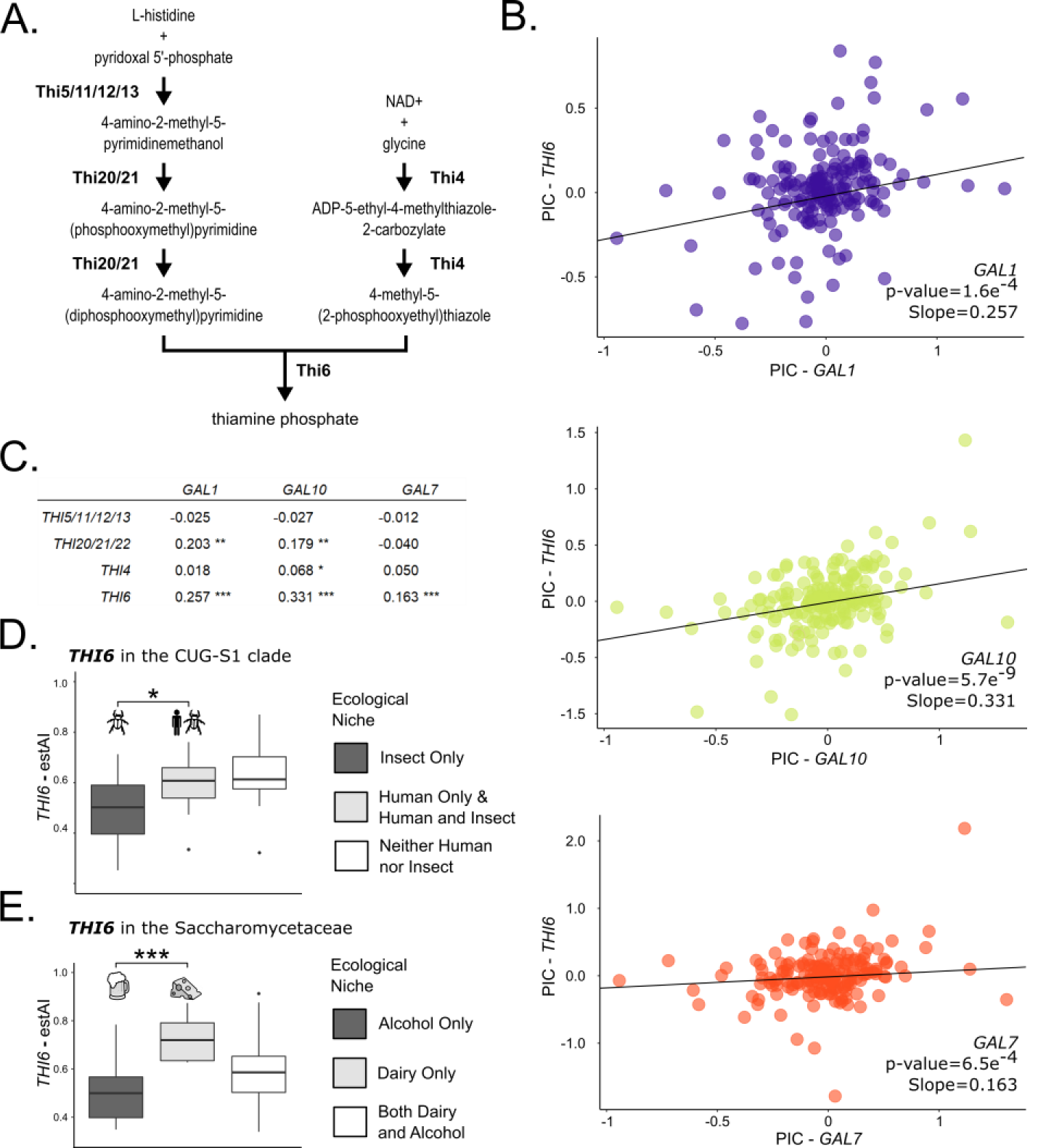
Codon optimization in the *GAL* pathway is positively and significantly correlated with optimization of multiple thiamine biosynthesis proteins. A) The two branches of the thiamine biosynthesis pathways present in budding yeasts converge on *THI6*. B) The PIC correlation between codon optimization in the *GAL* genes and *THI6* in species with evidence of genome-wide translational selection on codon usage (s-value >=0.5). The strongest correlation is between *GAL10* and *THI6* (182 species), followed by *GAL1* (168 species), and *GAL7* (170 species). C) Optimization in *GAL10* is also correlated with optimization of the *THI20*/*THI21/THI22* gene family and *THI4*– these genes encode the enzymes upstream of *THI6*. Optimization in *GAL1* is additionally correlated with *THI20*/*THI21/THI22* optimization. There is no correlation between optimization in the *THI5/THI11/THI12/THI13* gene family and any *GAL* genes. D) Optimization in *THI6* is significantly greater in CUG-Ser1 clade species associated with human or human and insect ecological niches (14 species) when compared to species associated only with insect ecological niches (48 species) (p-value=0.011). Twelve species were not associated with either human or insect ecological niches. E) Optimization in *THI6* is significantly higher in Saccharomycetaceae species associated with dairy ecological niches (6 species) versus those associated with alcohol ecological niches (16 species) (p-value=1.9e^-4^). Ten species with *THI6* are associated with both ecological niches, and 33 species are not associated with either environment.

### *GAL* codon optimization is associated with specific ecological niches

We further hypothesized that adaptation to specific ecological niches is associated with increased expression of the *GAL* pathway. Based on preliminary tests across 50 previously characterized ecological niches [54,60] for 114 species, we conducted an extensive literature search for the four ecological niches of interest—dairy, alcohol, human, insect—to maximize the number of species with ecological information. We uncovered two examples of niche-specific codon optimization (Figure 3): in the CUG-Ser1 clade, we found that *GAL* gene optimization was significantly higher in species that have been isolated from human-associated ecological niches versus those that have been isolated from insect-associated niches; and in the Saccharomycetaceae, we found *GAL* gene optimization was significantly higher in species isolated only from dairy-associated niches compared to species isolated only from alcohol-associated niches.

*CUG-Ser1 clade:* Among CUG-Ser1 species that exhibit high genome-wide evidence of translational selection on codon usage (s-value ≥ 0.5), we found that *GAL* gene optimization was significantly higher (p<0.05) in species from human-associated ecological niches or human-and insect-associated niches versus those that have been isolated from insect-associated niches only (57 species; Figure 3A). Only two species were found in human-associated niches and not insect-associated niches, *Debaryomyces subglobosus* and *Cephaloascus fragrans*; thus, we combined the human-associated species with the human-and insect-associated species into one group for subsequent analyses. Recent work has shown that many opportunistically pathogenic budding yeasts are likely to be associated with both environmental and human niches [94]. The 13 CUG-Ser1 species isolated from humans with genome-wide evidence of selection on codon usage had a mean optimization of 0.74, 0.76, and 0.69 for *GAL1, GAL10*, and *GAL7*, respectively.

We also found that *GAL1, GAL10*, and *GAL7* optimization was significantly higher (Wilcoxon rank sum test p-values of 0.035, 0.014 and 0.003, respectively) in species from human-associated ecological niches than insect-associated niches only, irrespective of genome-wide evidence of translational selection (88 species). For example, the major human pathogen *Candida albicans* does not have genome-wide evidence for high levels of translational selection but has a very high *GAL10* codon optimization (estAI = 0.86). While *C. albicans* may not have evidence of genome-wide selection on codon optimization, a previous analysis suggests that at least 17% of genes in the *C. albicans* genome have likely experienced selection on codon usage [59].

Other opportunistic human pathogenic species with very high *GAL10* codon optimization (estAI > 0.8) include *Candida dubliniensis* [95], *Meyerozyma caribbica* [96], *Candida tropicalis* [97], *Meyerozyma guilliermondii* [98], and *Clavispora lusitaniae* [99]. The optimization of *GAL10* in human pathogenic species is consistent with findings that *GAL10* expression is upregulated during *C. albicans* growth in the mammalian intestinal track [100]. Furthermore, *GAL10* in *C. albicans* is required for cell-wall integrity, resistance to oxidative stress, and other virulence-related traits, even in the absence of galactose [85]. This suggests that *GAL10* may play an additional role, outside of galactose metabolism, in the CUG-Ser1 clade.

Interestingly, the highest *GAL10* optimization (average estAI = 0.93) in the CUG-Ser1 clade is found in *Spathaspora* species. While many *Spathaspora* species have been isolated from insects, four of the five species studied here *(Sp. girioi, Sp. hagerdaliae, Sp. gorwiae*, and *Sp. arborariae*) have been isolated only from rotting wood [101,102]. This observation is particularly interesting given the hypothesis that some features of saprophytic fungi, such as *Aspergillus fumigatus* and *Cryptococcus* spp., enable or predispose them to colonize human hosts [103,104]. Moreover, some pathogenic budding yeasts, including *C. albicans* and *C. tropicalis*, have recently been associated with soil [94].

*Saccharomycetaceae*: Among Saccharomycetaceae species, we found that *GAL* optimization is significantly higher (p<0.05) in those that have been isolated only from dairy-associated niches compared to species isolated only from alcohol-associated niches (19 species; Figure 3B.) Only one species isolated from either dairy or alcohol, namely the alcohol-associated *Lachancea thermotolerans*, did not have evidence of genome-wide translational selection on codon usage. The four species isolated only from dairy-associated niches (*Kluyveromyces lactis, Naumovozyma dairenensis, Vanderwaltozyma polyspora*, and *Kazachstania turicensis*) have mean codon optimization values of 0.90, 0.88, and 0.84 for *GAL1, GAL10*, and *GAL7*, respectively. The ten species that are only from alcohol-associated niches (Supplementary table 2) have mean codon optimization values of 0.73, 0.61, and 0.59 for *GAL1, GAL10*, and *GAL7*, respectively. In many dairy environments, there are large microbial communities that often consist of lactic acid bacteria that convert lactose into glucose and galactose, which can subsequently be used in the *GAL* pathway [105,106]. The natural presence of galactose in dairy-associated environments is the likely driver of *GAL* codon optimization.

Species found in both dairy-and alcohol-associated niches have a range of optimization values that generally encompasses the values observed for species from dairy-or alcohol-only niches. It is likely that this group (associated with both dairy and alcohol niches) contains species or populations that are better adapted to one niche than the other. It is not possible, however, based on current literature to disentangle these two categories. For example, the species *Kluyveromyces marxianus* has been isolated from chica beer [107], cider [108], kombucha [109], and mezcal liquor [110]. However, *K. marxianus* is a well-known “dairy-yeast” frequently found in both natural [111,112] and industrial dairy products [113]. Codon optimization of the *GAL* enzymatic pathway is also very high in *K. marxianus* with an average estAI of 0.92. We hypothesize that the high *GAL* codon optimization in *K. marxianus* is a result of its association with dairy and with the ability of *K. marxianus* to metabolize lactose into glucose and galactose [44]. There are two species that are associated with both dairy and alcohol niches whose *GAL* codon optimization values are higher than the maximum value observed in alcohol-only species— *Naumovozyma castellii* and *Kazachstania unispora*. Based on this we hypothesize that these species are well adapted to dairy-associated environments.

### Differential *GAL* pathway optimization in *Kluyveromyces*

The genus *Kluyveromyces* provides an example of how codon optimization varies between closely related species that differ in their ecological niches (Figure 4). Two of the four species in this clade have not been isolated from either dairy or alcohol; *Kluyveromyces aestuarii* has been isolated from marine mud and seawater while *Kluyveromyces dobzhanskii* has been isolated from flies, plants, and mushrooms [60]. Of the four species represented here, only *K. dobzhanskii* is not known to metabolize lactose into glucose and galactose [60]. While all four species are capable of growing on galactose, *GAL* gene codon optimization is much higher in the two species with dairy-associated ecological niches, *Kluyveromyces lactis* and *Kluyveromyces marxianus* (Figure 4A.). Codon optimization for *GAL* genes is greater than 75% of the genome (estAI > 0.75) for *K. lactis* and *K. marxianus*. In *K. marxianus*, the optimization of *GAL1* and *GAL10* (estAI 0.93 and 0.94) is nearly that of the average ribosomal gene (estAI 0.99; Figure 4B). Ribosomal genes, which are among the most highly expressed genes in the genome, are known to be highly optimized in a broad range of species [114]. In contrast, optimization values for *GAL* genes in *K. aestuarii* and *K. dobzhanskii* are nearer to the mean (mean estAI values of 0.63 and 0.46, respectively; Figure 4B).

We hypothesized that the low *GAL* optimization in *K. aestuarii* and *K. dobzhanskii* was due to a relaxation in translational selection on the *GAL* pathway. To test this hypothesis, we estimated the rate of synonymous site evolution using PAML in the *GAL* genes and *VPS4*, a randomly chosen KEGG ortholog annotated in all 4 species. In each of the *GAL* genes, the branch length for *K. aestuarii* was the longest, and is at least double in length relative to the other branches in the *GAL7* and *GAL10* gene trees (Figure 4C; Supplementary Figure 7). The branch lengths of *K. dobzhanskii* were similar to those of *K. marxianus* in the trees of all three *GAL* genes. This pattern was not seen in the randomly chosen *VPS4* gene. This result suggests that relaxed selection on the *GAL* genes may exist in *K. aestuarii*, but not *K. dobzhanskii*, or that the relaxation may have persisted longer in *K. aestuarii*. Increased sampling in this clade would improve our understanding of the selective forces at work.

### *GAL* optimization is correlated with optimization in the thiamine biosynthesis pathway

In general, multiple metabolic pathways, as opposed to a single one, likely contribute to adaptation to an ecological niche [54,115]. To identify additional pathways associated with galactose optimization, we tested whether levels of codon optimization in *GAL* genes were significantly correlated with levels of codon optimization in other KEGG orthologs (KOs). We identified 78 / 2,572 KOs with a significant positive or negative association with *GAL* optimization (PIC, multiple test corrected p-value <0.05; Supplementary Table 3). One of the strongest positive associations (8^th^ smallest p-value in *GAL10* out of 28 KOs with significant positive associations) was with *THI6* (KO K14154), a member of the thiamine biosynthesis pathway (Figure 5A and B). We expanded our analysis to the two branches of the thiamine biosynthesis pathway present in the budding yeast subphylum that converge on *THI6* (Figure 5C). On the branch of the thiamine biosynthetic pathway that begins with the substrates pyridoxal 5’phosphate and L-histidine, we found significantly (p<0.05) correlated codon optimizations between the *THI20/THI21/THI22* gene family and the *GAL* genes *GAL1* and *GAL10* (Figure 5C). In the other branch of the pathway, codon optimization in *THI4* is only correlated with *GAL10* (Figure 5C). Among genes involved in thiamine biosynthesis, the strongest association with the *GAL* pathway was seen in *THI6* where there was a significant positive association with optimization in all three *GAL* genes (Figure 5B). The positive correlation seen using PIC suggests that this association does not reflect phylogenetic constraint but adaptation.

Support for the notion that ecological adaptation explains the correlation between the thiamine biosynthesis and *GAL* pathways can be found in both major clades examined. Within the CUG-Ser1 clade, there is a significantly higher (p<0.05) *THI6* codon optimization in species associated with either human or insect ecological niches when compared to species only isolated from insect ecological niches (Figure 5D). The difference in *THI6* codon optimization is even more significant (p<0.001) in the Saccharomycetaceae where *THI6* codon optimization is higher in species only associated with dairy ecological niches and not alcohol ecological niches (Figure 5E). Many lactic acid bacteria found in dairy environments, such as *Lactobacillus brevis*, require extracellular thiamine [116]. One possible model is that, in dairy communities containing lactic acid bacteria and yeasts, stiff extracellular competition for thiamine may boost the expression of thiamine biosynthesis genes in these yeasts. Alternatively, the thiamine biosynthesis and galactose metabolism pathways may be connected by metabolic intermediates [117]. It is possible that both a biochemical and ecological explanation underlie the correlation between codon optimization in the *GAL* and thiamine biosynthesis pathways.

## CONCLUSIONS

Here we use reverse ecology to connect genotype (codon optimization) with phenotype (growth rate on galactose) and ecology (isolation environment) across an entire evolutionary lineage (budding yeasts). By studying a well-known metabolic pathway in a diverse microbial subphylum, we provide a proof of concept for the utility of codon optimization as a genomic feature for reverse ecology. Our discovery of optimization in the *GAL* pathway in dairy-associated Saccharomycetaceae and human-associated CUG-Ser1 yeasts is consistent with the known functional roles of the enzymes in the pathway. The complete *GAL* pathway metabolizes lactose, a component of dairy environments, into usable energy [118]. The *GAL10* gene is associated with phenotypes associated with human colonization in CUG-Ser1 yeasts [85]. Similarly, in the *Kluyveromyces* species found on dairy-associated niches that are able to metabolize lactose into glucose and galactose, there is high optimization in this pathway compared to closely related species not associated with dairy. Interestingly, examination of codon optimization in the gene sets of the four *Kluyveromyces* species studied here would have identified at least *K. marxianus* as a potential dairy-associated yeast, even in the absence of any knowledge about its isolation environments. Thus, genome-wide examination of codon optimization in fungal, and more generally microbial, species can generate specific hypotheses about metabolic ecology in species for which ecological data are lacking. These results are especially promising as this method can be applied directly to genomic data—which is the only source of information for microbial dark matter known only from DNA [119]. Finally, using an unbiased approach, we identified a strong correlation between optimization in the thiamine biosynthesis pathway and the *GAL* pathway. This novel finding suggests that codon optimization may also be useful for identifying co-regulated or correlated pathways in microbial, including fungal, species.

## Acknowledgements

We thank the members of the Rokas and Hittinger labs for helpful discussions.

## Data availability

All analyses were done on publicly available and published genome assemblies and annotations. The codon optimization values were obtained from the figshare repository from LaBella et al. 2019 (https://doi.org/10.6084/m9.figshare.c.4498292.v1). Additional sequence data generated in this project, including the reference and annotated gene sequences, are stored in the figshare repository associated with this manuscript and will be made publicly available upon acceptance to a peer-reviewed journal. Reviewers can access the figshare repository through the private link:. All other information and data generated are available in the supplementary files.

## Notes

### Competing Interest Statement

The authors have declared no competing interest.

