## Supplementary Tables and Figures for "Signatures of optimal codon usage predict metabolic ecology in budding yeasts": LaBella_et_al_2020_supplementary_figures.pdf

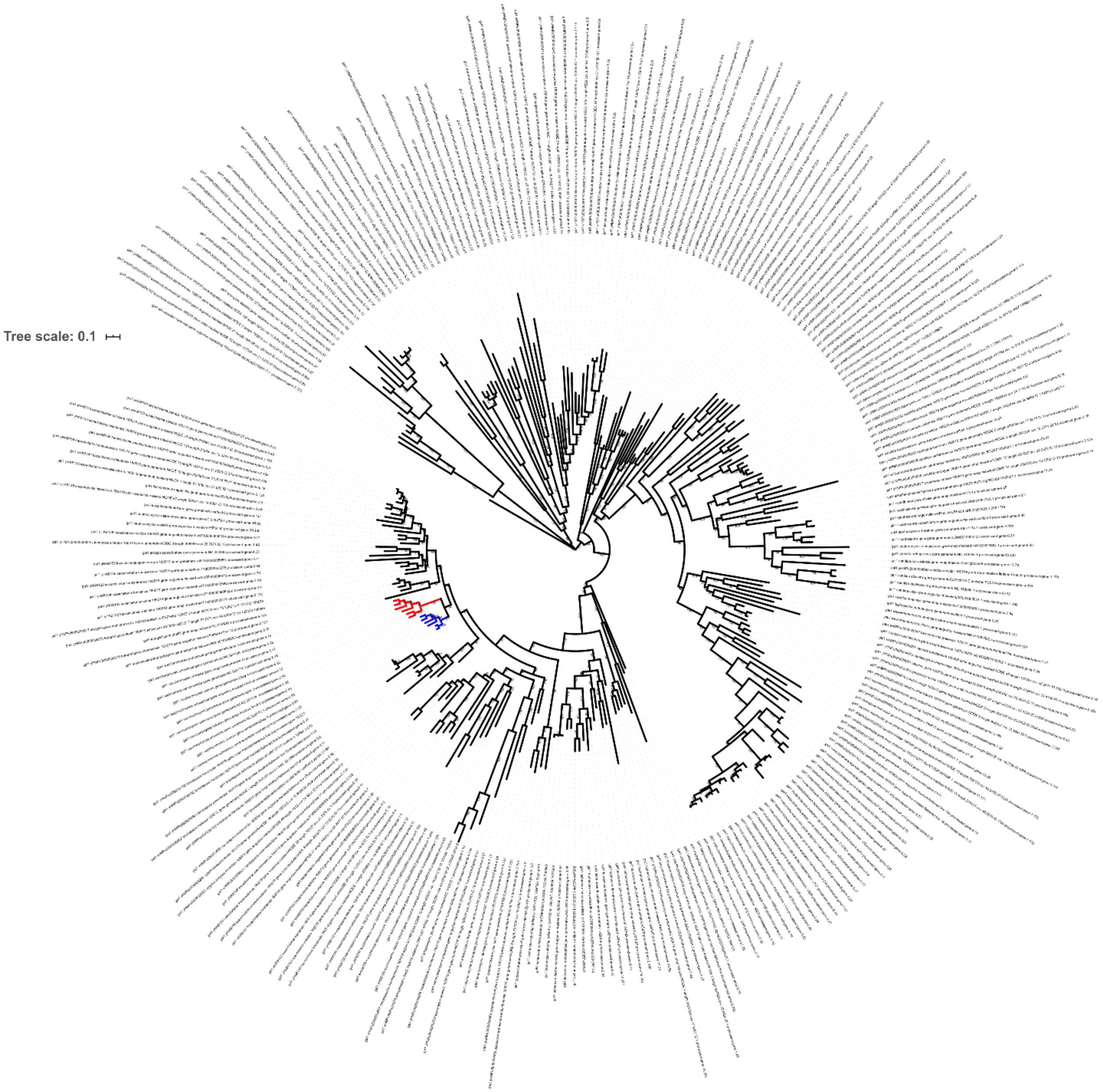


**Supplementary Figure 1: ML tree of Gal1 protein sequences for the Saccharomcotina**. The red and blue clades represent the *Saccharomyces* species known to contain Gal3. The red sequences were identified as Gal3 based on the placement of the known *Saccharomyces cerevisiae* Gal3 sequence.

A.

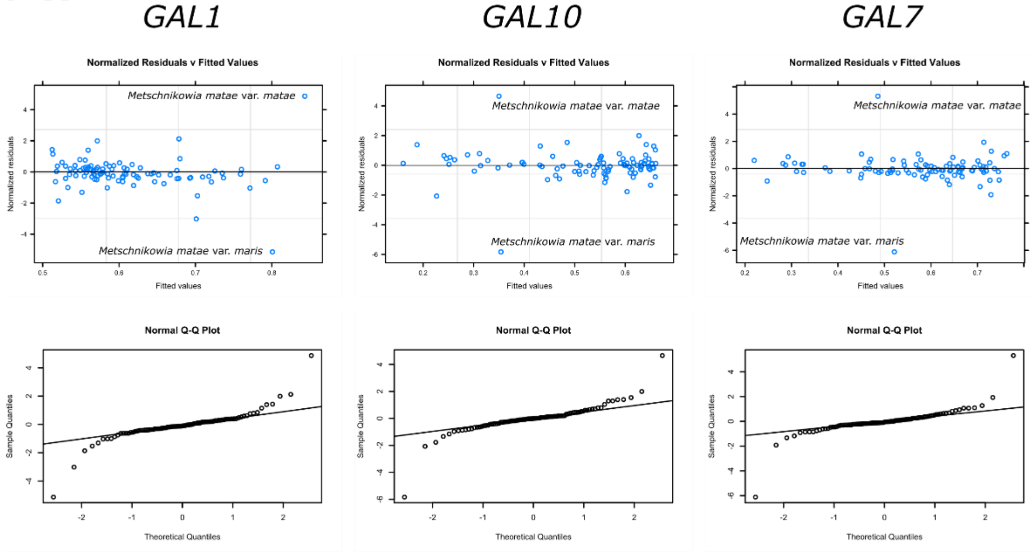


B.

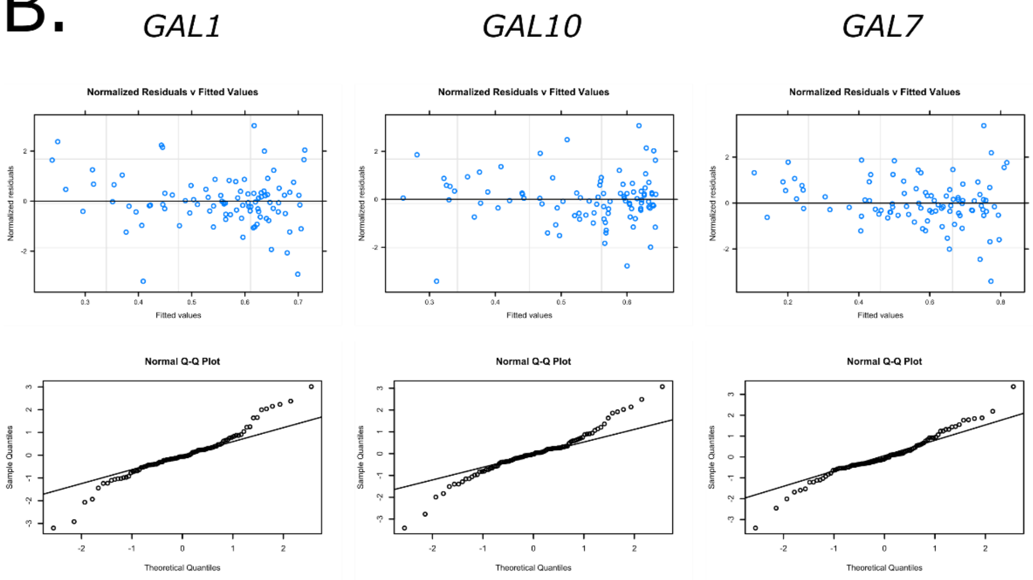


**Supplementary Figure 2: Residual and Q-Q normalized plots of phylogenetic generalized least squares (PGLS) analysis of codon optimization and normalized growth-rate on galactose containing medium.** A. The residual versus fitted analysis shows two outlier strains: *Metschnikowia matae* var. *matae* and *Metschnikowia matae* var. *maris.* B. Residual and Q-Q normalized plots after removal of *Metschnikowia matae* var. *matae*. No clear outliers remained after removal of this species.

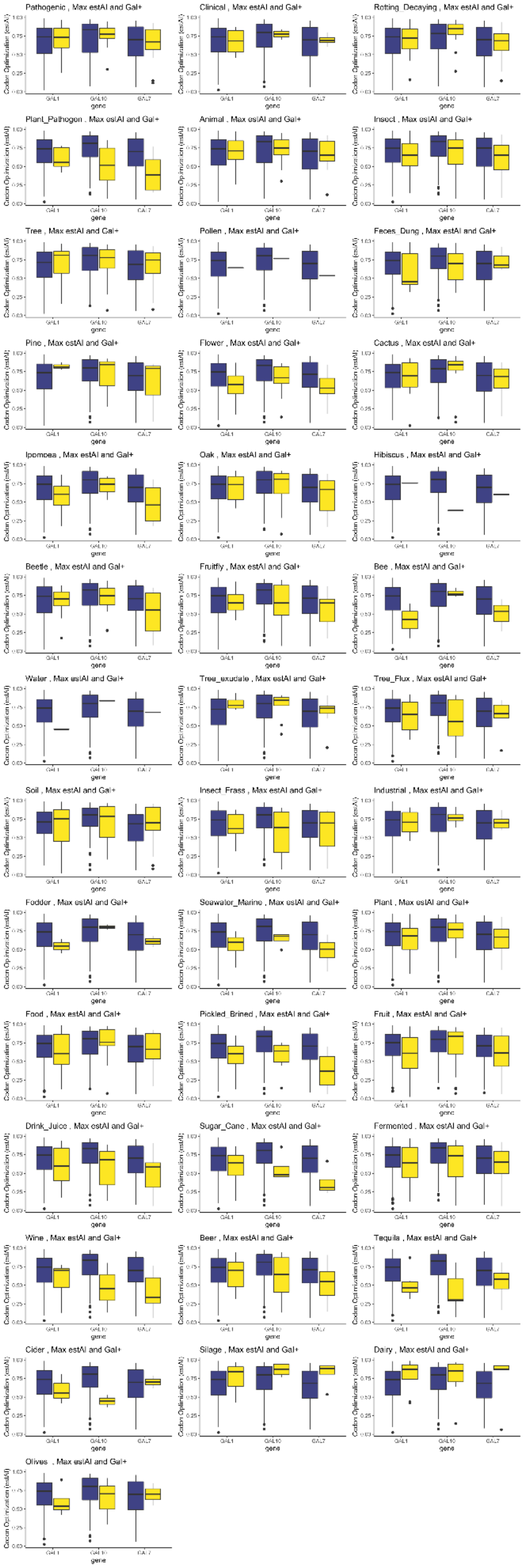


**Supplementary Figure 3: Initial analysis of *GAL* codon optimization in species isolated from particular ecological niches versus those that have not been isolated from a niche.** Blue bars are the codon optimization values for species that have not been isolated from the particular ecology. Yellow bars are the codon optimization values for species that have been isolated from that ecology. Ecological information was tested in 50 isolation environments from data collated from *The Yeasts: A Taxonomic Study* as recorded by Opulente et al. 2018.

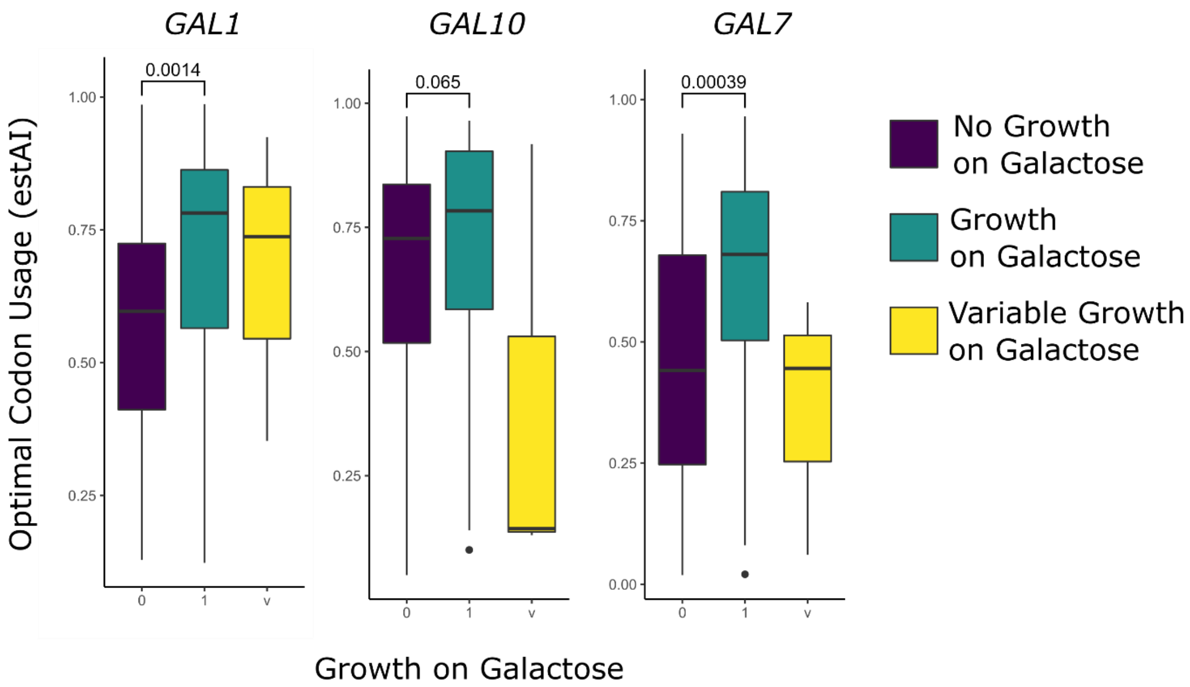


**Supplementary Figure 4: Wilcoxon rank sum test of *GAL* codon optimization versus binary data for growth on galactose.** A total of 170 species were included in this analysis.

### A. Saccharomycetacea

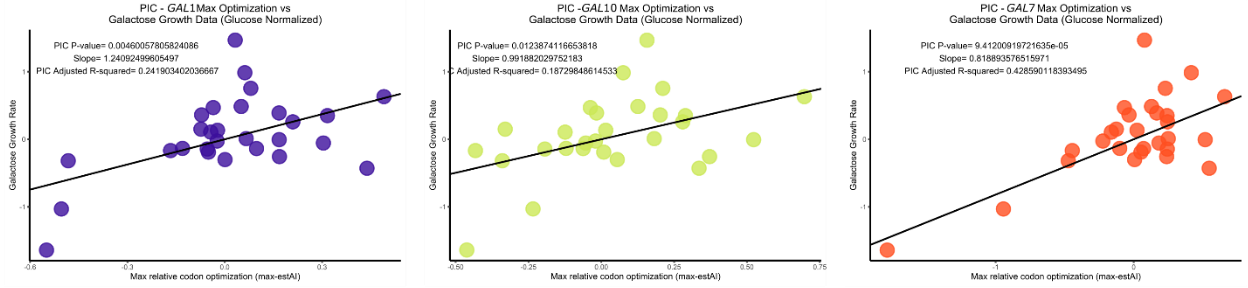


### B. CUG-Ser1 Clade

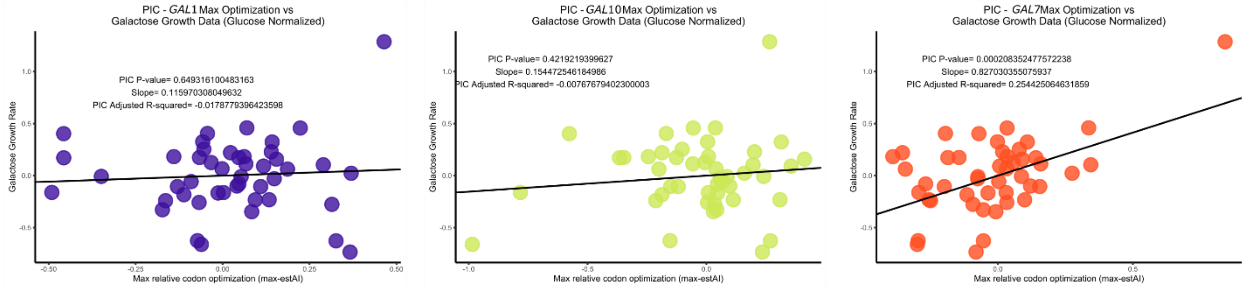


**Supplementary Figure 5: PIC analysis of *GAL* codon optimization and growth rate on galactose containing medium (normalized to growth rate on glucose containing media).** A) The phylogenetically corrected correlation between *GAL* codon optimization and quantitative growth on galactose containing medium is significant in all genes when only species from the family Saccharomycetacea are considered. 29 species were included in this analysis. B) The phylogenetically corrected correlation between *GAL* codon optimization and quantitative growth on galactose containing media is only significant in *GAL7* when only the CUG-Ser1 major clade species are considered. This analysis includes 47 species.

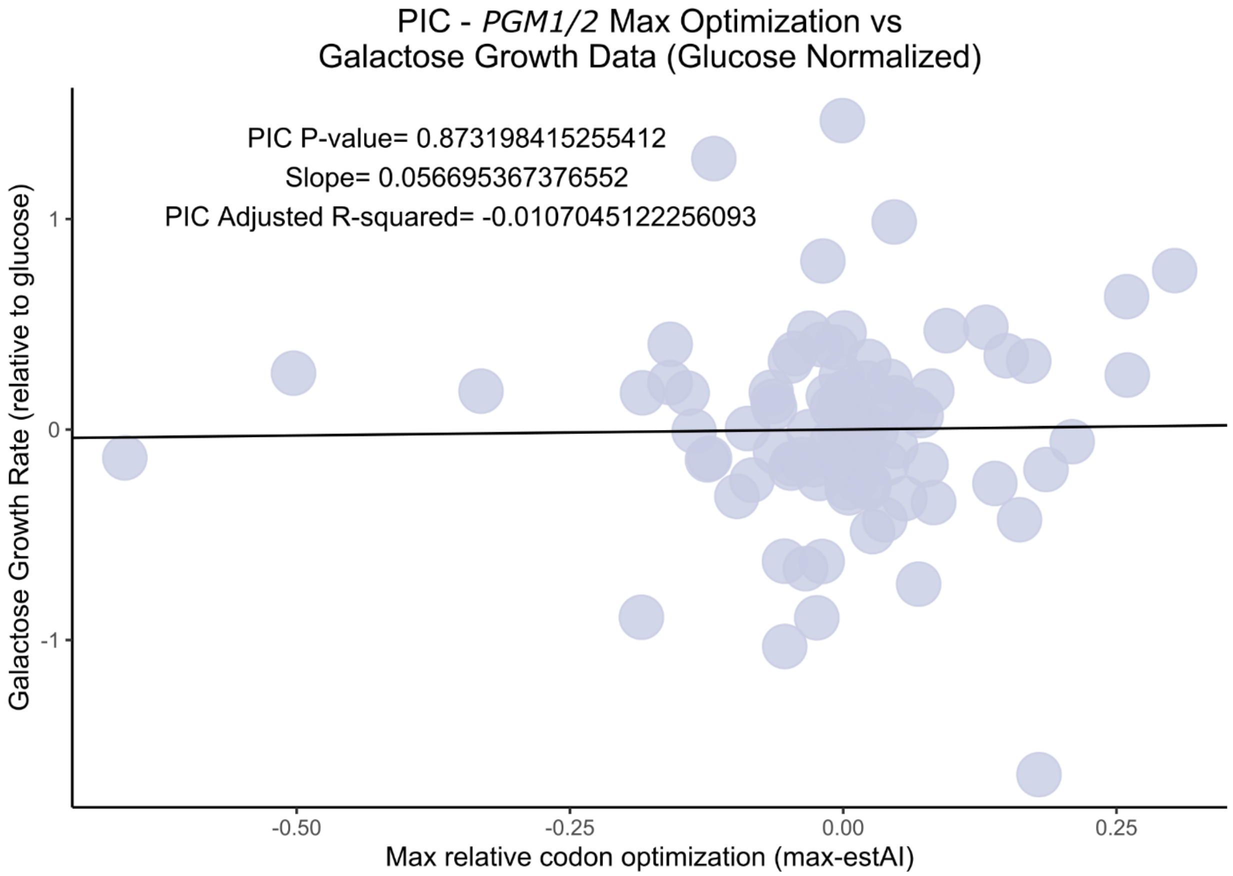


**Supplementary Figure 6: PIC analysis of correlation between growth-rate on galactose containing medium and codon optimization in the gene *PGM1/2*.** This analysis suggests that optimization in *PGM1/2* does not contribute to growth on galactose containing medium.

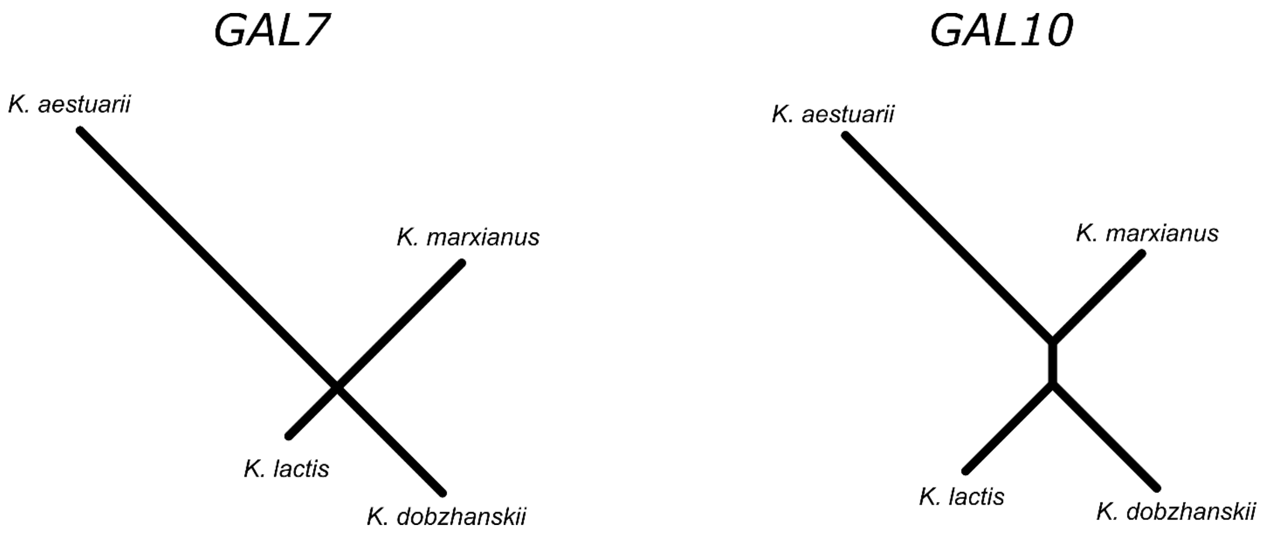


**Supplementary Figure 7: The unrooted trees show the estimated rate of synonymous substitutions in the *GAL7* and *GAL10* genes for the *Kluyveromyces* species.** The internal branch of *GAL7* is 0.00027.
